# Mapping Gene Drive Dynamics onto Mendelian Models

**DOI:** 10.64898/2026.01.28.702305

**Authors:** Qimo Wan, Zihang Wen, Gili Greenbaum, Oana Carja

## Abstract

CRISPR-based gene drives bias their own transmission and can spread even when deleterious, giving rise to evolutionary dynamics that can be substantially more complex than those governed by standard Mendelian inheritance. Identifying conditions under which gene-drive dynamics can be faithfully approximated by Mendelian models would therefore enable the extensive theoretical toolkit of classical population genetics to be applied to gene-drive systems. Here, we develop a general mapping framework that translates gene-drive models into dynamically equivalent Mendelian models, allowing their behavior to be analyzed using classical theory. By deriving both haploid and diploid effective-parameter mappings, we identify Mendelian models that closely reproduce allele-frequency trajectories of gene drives across a wide range of conversion rates, fitness costs, and dominance effects. We delineate the regions of the parameter space where a one-parameter haploid approximation provides an accurate first-order representation, and where incorporating dominance in a diploid mapping substantially improves fidelity and recovers internal equilibria and threshold behavior. Analytic approximations yield efficient mappings across most of the drive parameter space, while a trajectory-based grid search further improves accuracy near nonlinear regime boundaries. To demonstrate the utility of this framework, we apply it to predicting gene swamping in a two-deme migration-selection model and show that the mapped Mendelian system accurately forecasts transitions between fixation and loss under three relevant release scenarios: environmental variation in fitness, engineered fitness asymmetries, and environment-dependent conversion. Together, these results establish a theoretical bridge between non-Mendelian gene drives and classical population genetic models, providing an interpretable and computationally efficient foundation for predicting gene-drive outcomes and guiding the design of gene drive systems and deployment strategies.

## Introduction

Gene drives are genetic elements that bias their own transmission, allowing them to spread rapidly through populations even when their carriers have reduced fitness (5, 12). Unlike Mendelian inheritance, in which alleles have a 50% probability of transmission to offspring, gene drives can achieve substantially higher transmission probabilities (1, 3,4,11,24,32, 35). The advent of CRISPR-based “homing” drives (17, 20) has transformed gene drives from a largely theoretical concept into a practical genetic engineering strategy, raising the prospect of unprecedented control over the spread of engineered traits in wild populations (6, 12). As a result, gene drives have emerged as powerful candidate tools for applications in public health, agriculture, and conservation biology, including the control of vector-borne diseases and invasive species (15, 19).

Because releasing gene drives into natural populations carries substantial ecological and ethical risks, mathematical and computational models have become indispensable for characterizing their spread, stability, and potential consequences prior to any field deployment (10, 16, 18, 21, 22, 25, 26, 33). These models provide a critical framework for evaluating both the efficacy of gene-drive systems and the conditions under which they can be contained across a wide range of ecological and evolutionary settings. However, gene-drive dynamics are far less tractable than those governed by classical Mendelian inheritance. Biased gene conversion fundamentally alters allele transmission, introducing strong nonlinearities and frequency-dependent effects that complicate analytical treatment and obscure intuitive interpretation (29, 33).

As a consequence, many gene-drive models lack closed-form solutions and resist direct comparison across different drive architectures or parameter regimes, forcing researchers to rely heavily on numerical simulations tailored to specific designs and ecological contexts. While such simulations are indispensable, their specificity can make it difficult to synthesize results across studies or to extract general, interpretable principles. These limitations point to the need for theoretical frameworks that connect gene-drive dynamics to simpler, classical population-genetic models: frameworks that preserve biological realism while enabling broader insight and comparison.

In this study, we introduce a mapping framework that formally connects non-Mendelian gene-drive systems to classical Mendelian population-genetic models. By deriving haploid and diploid effective-parameter mappings, we ask under what conditions three core gene-drive parameters—fitness cost, dominance, and conversion efficiency—can be translated into equivalent Mendelian selection parameters, and when such reductions break down. This approach allows complex gene-drive dynamics to be approximated by simpler, well-studied Mendelian models, providing an interpretable and computationally efficient framework while explicitly delineating the limits of Mendelian representations.

As an application of this framework, we analyze gene swamping in gene-drive systems. Gene swamping occurs when sufficiently high migration rates prevent a gene drive from establishing and ultimately purge it from the population, a behavior that has been proposed to be useful in the design of deployment programs (21). Using a classical two-deme migration-selection model (8, 9, 28, 36), we show that the Mendelian mapping accurately predicts gene-drive outcomes across a broad range of ecological and engineered scenarios, including spatial variation in fitness, engineered habitat-specific fitness asymmetries, and environment-dependent conversion efficiencies. In each case, the mapped Mendelian model correctly identifies the conditions under which swamping occurs, demonstrating the framework’s ability to understand containment and reversal dynamics in spatially structured populations.

Taken together, our results establish a theoretical bridge between non-Mendelian gene drives and classical population genetics. By translating gene-drive dynamics into effective Mendelian parameters, this framework brings conceptual clarity to otherwise complex dynamics, offers substantial computational efficiency, and opens access to a century of population-genetic theory. More broadly, it enables gene-drive behavior to be approximated and interpreted using familiar evolutionary principles.

## A framework for mapping gene drive models onto Mendelian models

### Modeling gene drive dynamics

For the models describing the behavior of gene-drive alleles, we consider the well-studied CRISPR-based homing drive model (18, 33), in which the gene drive construct can convert heterozygotes into gene-drive homozygotes. In this model, the frequency of the gene-drive allele is tracked using a recursion equation, with dynamics governed by three parameters: the fitness cost of the gene-drive allele s, the dominance of the gene-drive allele relative to the wild-type allele h, and the probability c that heterozygotes are converted into gene-drive homozygotes (**Figure 1A**). Although gene-drive conversion can occur either prior to viability selection (zygotic conversion) or after viability selection in the germline (10, 16, 18), here we consider germline conversion, since this assumption aligns more closely with empirical observations (12, 23, 27) and most recent modeling studies.

**Figure 1.**
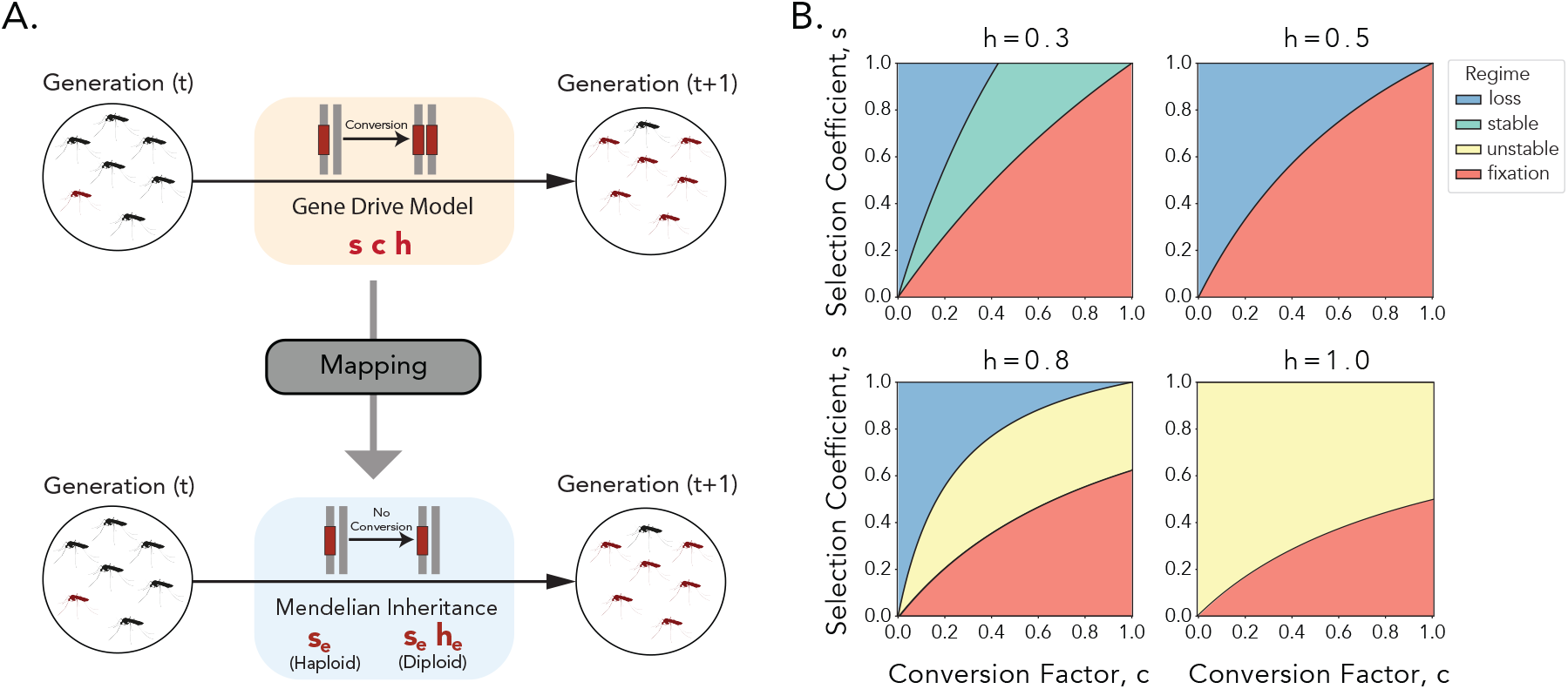
A framework for mapping gene drive models onto Mendelian models. **A.** Our goal is to develop a general framework that maps gene-drive models onto the Mendelian model that best reproduces the allele-frequency trajectory of the gene-drive model. We therefore look for a mapping of the gene drive model with three parameters (s, c, h) that produce similar changes in frequency over a generation, either in a haploid Mendelian model with a single parameter s_e_ or a diploid Mendelian model with two parameters (s_e_, h_e_). **B**. Across the parameter space defined by (s, c, h), the gene drive model can generate up to four qualitatively distinct outcomes: fixation, loss, a stable internal equilibrium, and an unstable internal equilibrium.

Under this model, the change in the frequency of the gene drive allele between generations t and (t + 1) can be formulated as

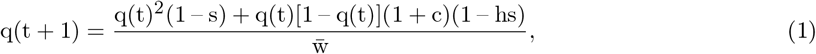

where 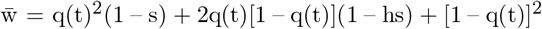 is the mean fitness of the population (16, 33). This formulation serves as our baseline model for gene-drive dynamics. Across the parameter space defined by (s, c, h), this system can generate four qualitatively distinct outcomes: fixation of the gene drive allele, loss of the gene drive allele, a stable polymorphic internal equilibrium, and an unstable internal equilibrium (**Figure 1B**)(16, 33).

### Mapping gene drive models onto Mendelian models

Our goal is to identify classical Mendelian models that reproduce the dynamics of a given gene-drive system. To achieve this, we develop a general framework that maps gene-drive models onto Mendelian models using either analytical approximations or computational searches.

We systematically map each gene-drive model onto one of two corresponding Mendelian model systems: a haploid Mendelian model or a diploid Mendelian model. This mapping allows us to determine which regions of the gene-drive parameter space, defined by the triplet (s, c, h), can be accurately approximated by a one-parameter haploid model or a two-parameter diploid model. In the haploid representation, the drive allele is treated as an allele with a relative selection coefficient s_e_, which we consider as the effective Mendelian selection coefficient corresponding to the gene drive model. Similarly, in the diploid representation, the corresponding Mendelian system is characterized by an effective selection coefficient s_e_ and an effective dominance coefficient h_e_. In other words, for each gene-drive parameter set (s, c, h), we identify the Mendelian parameter set, either (s_e_) or (s_e_, h_e_), that best reproduces the allele-frequency trajectory of the gene-drive model from one generation to the next (**Figure 1A**).

We implement two complementary mapping strategies: (i) an analytic approximation that yields a closed-form expression for the effective Mendelian coefficients, and (ii) a trajectory-matching grid-search procedure that identifies the Mendelian parameter values minimizing the mean squared error (MSE) between gene-drive and non–gene-drive allele-frequency trajectories over time (more details in the **Methods** section).

Because some of the Mendelian models capture only a subset of the long-term behaviors exhibited by gene-drive systems, for example, the haploid model permits only fixation or loss, we develop distinct mapping procedures tailored to each gene-drive outcome: fixation, loss, stable equilibrium, and unstable equilibrium. This structure enables systematic comparison between gene-drive and Mendelian models and establishes a unified framework for predicting gene-drive dynamics using classical population-genetic theory.

## Results

### Mapping onto Mendelian haploid model in the fixation regime

For gene-drive parameter combinations that fall within the fixation regime (red regions in **Figure 1B**), we begin by approximating the dynamics using a one-parameter haploid Mendelian model. In this simple model, the population consists of two alleles with relative fitnesses 1 and (1 – s_e_). To derive an analytical mapping between the gene drive and the Mendelian models, we follow the approach introduced in (33) and approximate the early dynamics of the gene drive by analyzing its behavior when the drive allele is rare. In this regime, drive homozygote frequencies are negligible (q^2^ ≈ 0), and the average fitness in the population is approximately one (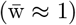). Under these assumptions, the haploid model recursion, 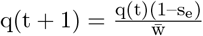,with 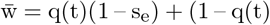 the mean fitness of the population, reduces to a linear approximation

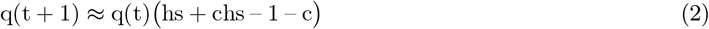

and we can define a mapping

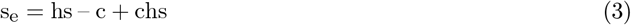

from the gene drive parameters (s, c, h) to an equivalent haploid Mendelian model with a single coefficient s_e_.

In addition to this analytical approximation approach, we also implemented a trajectory-matching grid-search procedure in which, for each gene drive configuration (s, c, h), we identify the corresponding value of s_e_ by performing a grid search over candidate values in the range 0.0 to 10.0. For each candidate non-gene-drive parameter, we compute the allele-frequency trajectories under both models and calculate the mean squared error (MSE) between the gene-drive and Mendelian trajectories over time. The value of s_e_ that minimizes the MSE is then recorded as the mapped parameter for that specific gene drive configuration.

Under both the analytic and grid-search approaches we observe similar mappings that produce negative effective selection coefficients s_e_ (i.e., a beneficial haploid allele)(**Figure 2**). In these mappings, for different c and h values, s_e_ increases approximately linearly with the gene-drive fitness cost s (**Figure 2A**). As expected, more deleterious gene drives map to more weakly selected, or nearly neutral, haploid dynamics. Across most of the fixation regime, both mapping approaches achieve consistently low mean-squared error (MSE), indicating that the haploid approximation accurately captures gene-drive fixation dynamics (**Figures 2B, C**). For gene-drive parameters that approach the boundaries separating fixation from other dynamical regimes, however, the haploid model becomes less able to reproduce the full dynamics (**Figure 2D**). In these transition regions the accuracy of the two approaches diverge: the analytic method exhibits a sharp increase in error, whereas the grid-search method maintains low error across the same parameter space (**Figures 2B, C**). This indicates that the grid-search approach is more robust to the nonlinearities and edge effects that arise near regime transitions, also when considering different dominance values, h (**Supplementary Figure S1**).

**Figure 2.**
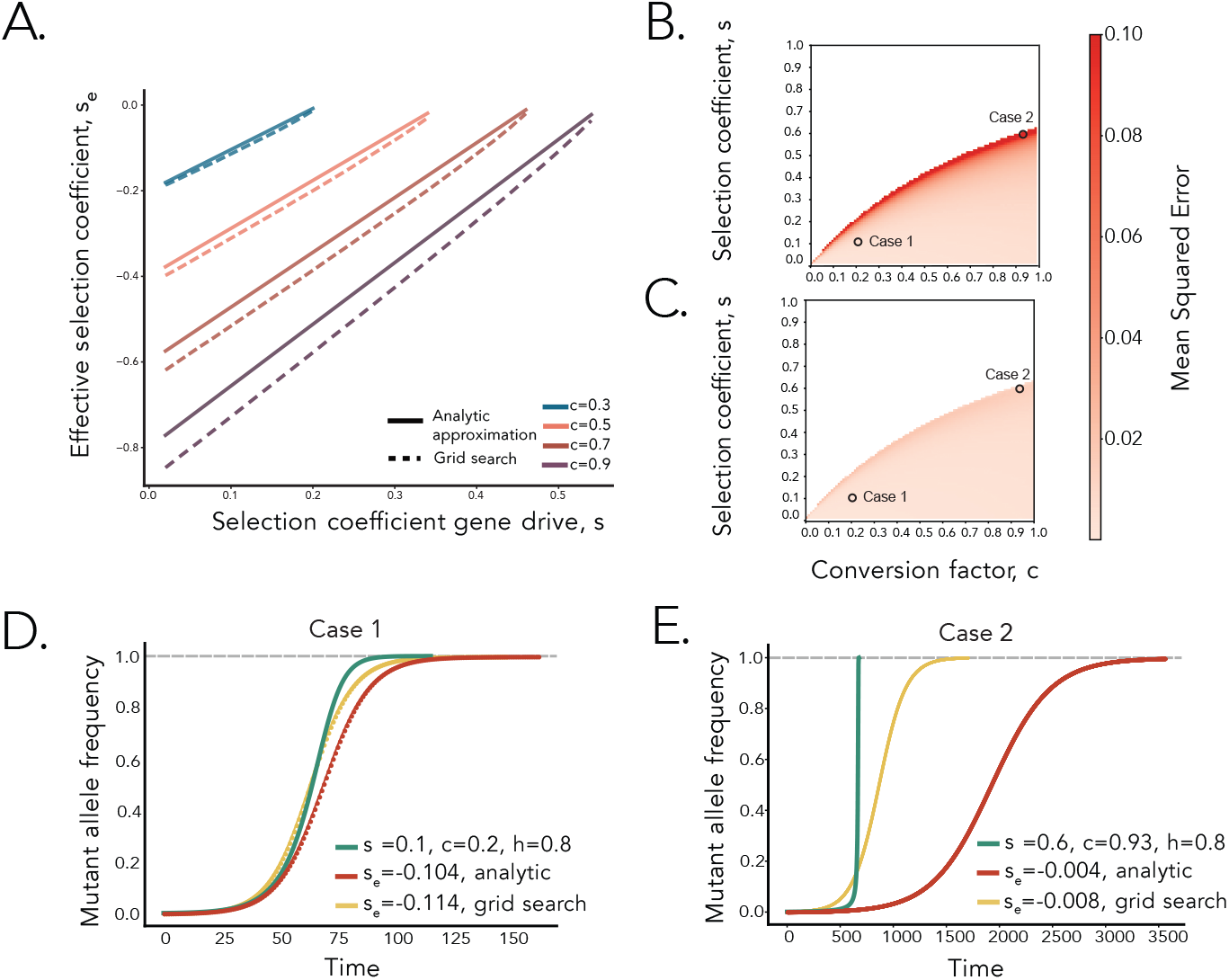
Fixation regime: Mapping to a haploid Mendelian model. **A.** Effective haploid selection coefficient, s_e_, as a function of the gene-drive selection coefficient, s, for h = 0.8 and different conversion rates, c. Solid lines denote analytic approximations using eq. (3), with dashed lines showing grid search results. **B, C**. Heatmaps of the mean squared error (MSE) between mapped haploid trajectories and the original gene-drive trajectories across (s, c) space, using the analytic mapping (**B**) and the grid-search mapping (**C**). **D, E**. Representative examples of gene-drive trajectories (green) and their mapped haploid approximations (analytic: red; grid search: yellow), for the two cases highlighted in panels **B** and **C**. All panels assume dominance h = 0.8 and initial gene-drive frequency q_init_ = 0.001.

Taken together, these results demonstrate that the haploid mapping provides an efficient and reliable approximation for most parameter combinations within the fixation regime, achieving high predictive accuracy with minimal computational cost. However, its declining performance near regime boundaries highlights the need to explore the accuracy of a two-parameter diploid mapping in regions where multiple equilibria or unstable threshold dynamics may arise.

### Mapping onto Mendelian diploid models in the fixation regime

We next consider a Mendelian diploid model that includes ordinary Mendelian inheritance with selection and dominance (s_e_, h_e_). In this framework, allele frequency dynamics follow the familiar recursion

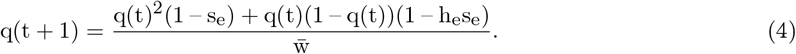

Here,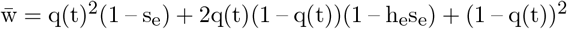 is the mean fitness of the diploid population.

To approximate gene drive dynamics using this Mendelian diploid model, we seek values of (s_e_, h_e_) such that, for a given allele frequency q, the change in frequency over one generation for the Mendelian (eq. (4)) model is similar to the gene drive (eq. (1)) model, leading to the condition:

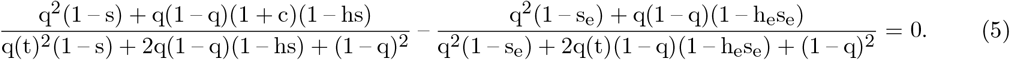

Because solving eq. (5) directly for a general q is analytically intractable, we approximate it using a Taylor expansion around q = 0, and obtain

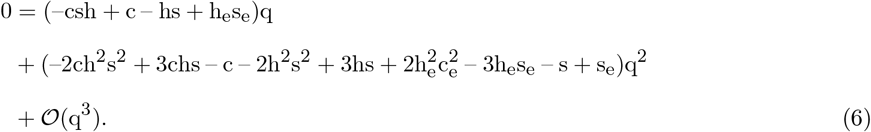

Imposing equality up to second order (i.e., setting the coefficients of q and q^2^ to zero) yields the system

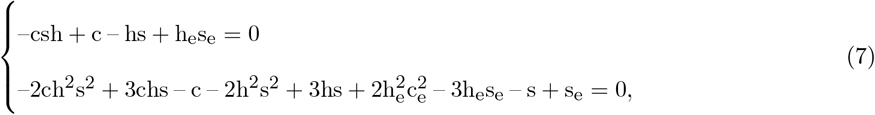

from which we obtain closed-form expressions for the effective parameters,

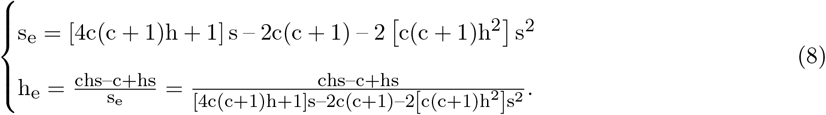

The effective selection coefficient s_e_ increases approximately linearly with s, when h or c are sufficiently small that higher-order (s^2^) terms can be further neglected, with a slope that depends on h. In contrast, the effective dominance parameter h_e_ varies nonlinearly with s, reflecting shifts of effective dominance rates under stronger selection. Corresponding grid-search results are shown in **Supplementary Figure S2**.

The mapping to the diploid Mendelian model, compared to the haploid Mendelian model, substantially improves the approximation to the gene drive model, because the introduction of the effective dominance parameter h_e_ captures heterozygote-specific effects. While the analytic mapping provides an efficient closed-form approximation that performs well across most of the fixation regime, the grid-search approach offers additional accuracy near nonlinear boundaries where dominance and conversion interact strongly (**Figures 3B, C**). Heatmaps across a wider range of dominance h values (**Supplementary Figure S3**) confirm that both analytic and grid-search diploid mappings remain accurate across dominance regimes.

**Figure 3.**
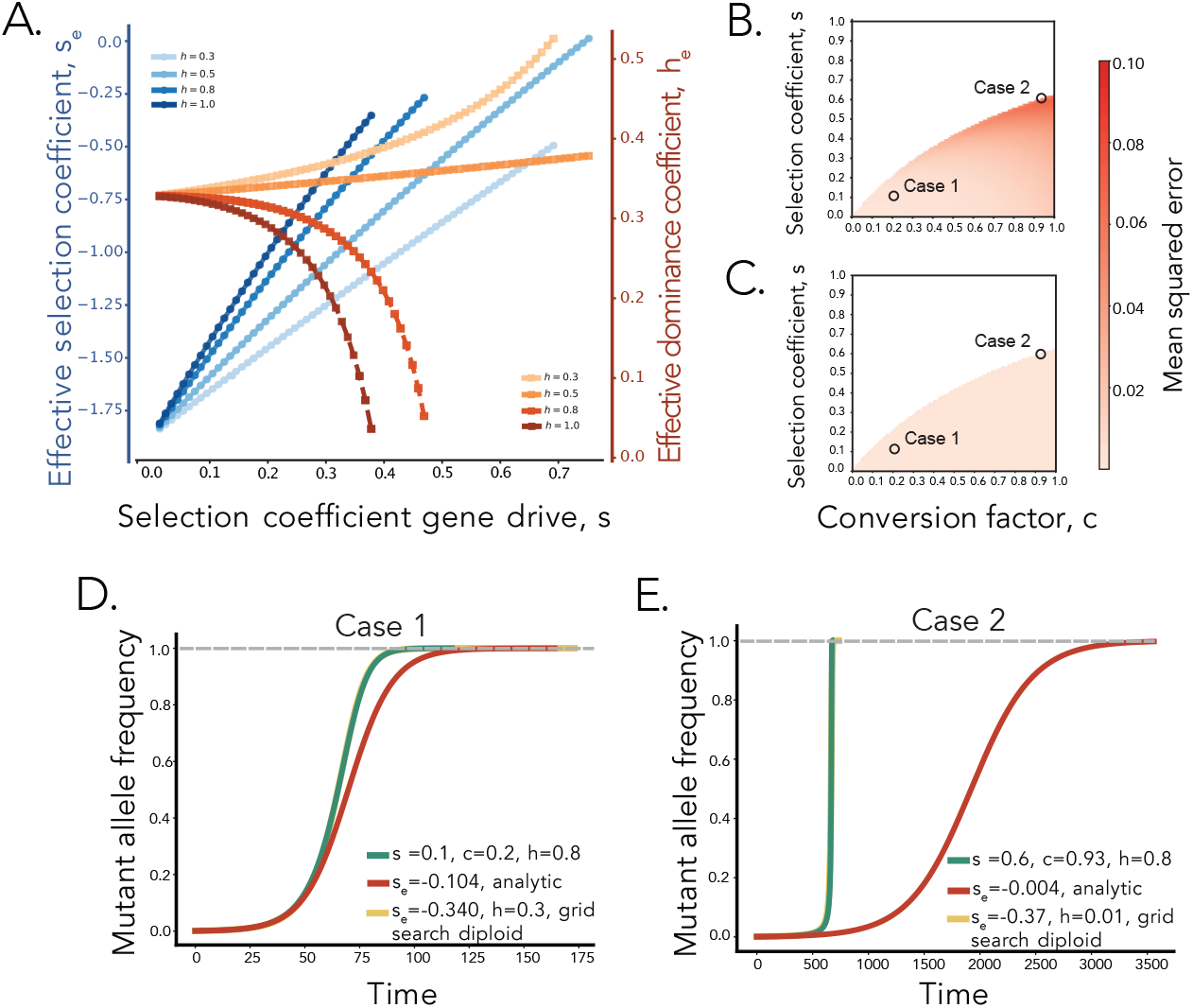
Fixation regime: Mapping to a diploid Mendelian model. **A.** Effective diploid selection coefficient (s_e_) and effective dominance coefficient (h_e_) as a function of the gene-drive selection coefficient (s), across different dominance coefficients (h), using eq. (8), with conversion rate c = 0.6. **B, C**. Heatmaps of the mean squared error (MSE) between mapped diploid trajectories and the original gene-drive trajectories across (s, c) space, using the analytic mapping (**B**) and the grid-search mapping (**C**). **D, E**. Representative examples of gene-drive trajectories (green) and their mapped haploid approximations (analytic: red; grid search: yellow), for the two cases highlighted in panels **B** and **C**. All panels assume initial gene-drive frequency q_init_ = 0.001.

Representative allele-frequency trajectories further highlight these trends (**Figures 3D, E**). For parameter combinations well within the fixation regime, both analytic and grid-search mappings closely track the gene drive allele frequency trajectory. Near regime boundaries, however, the analytic mapping underestimates the speed of increase and produces a slightly delayed fixation relative to the gene drive model. In contrast, the grid-search approximation continues to follow gene drive dynamics closely, reinforcing its robustness in parameter regions with strong nonlinearities.

### Mapping onto Mendelian diploid models in the unstable equilibrium regime

We next investigate the approximation and mapping procedure in the unstable-equilibrium regime. To approximate gene drive dynamics in this regime, we first require that both the gene-drive and diploid Mendelian models possess an internal equilibrium. Solving eq. (1) with q(t + 1) = q(t) yields the internal equilibrium 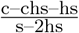 Similarly, solving eq. (4) for the effective selection and dominance parameters (s_e_, h_e_) provides the Mendelian model equilibria, with the internal equilibrium given by 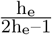.

Requiring these two internal equilibria to coincide, and matching the first-order Taylor expansions of the two models (see eq. (6)), yields

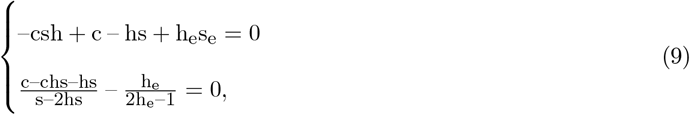

from which we obtain explicit expressions for the effective parameters

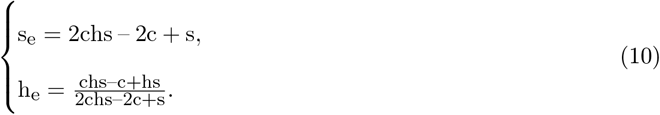

This result provides the analytical mapping between the gene drive and diploid Mendelian model for the unstable equilibrium regime.

For the grid search approach, because evolutionary outcomes in the unstable regime depend jointly on (i) the location of the internal equilibrium q_eq_ and (ii) the initial allele frequency q_init_, no single deterministic trajectory can capture the full range of possible dynamics. To obtain an accurate approximation using the grid-search procedure, for each candidate s_e_, we simulate trajectories in the Mendelian model across a range of initial allele frequencies q_init_∈{0, 0.005, 0.010, …, 1.0}, and compute the mean squared error (MSE) between the trajectories of the two models, averaged across all these q_init_ values.

The analytic mapping captures the overall linear relationship between s_e_ and s and performs well across most of parameter space (**Figures 4A, B**). Near the boundary separating the unstable and fixation regimes, however, particularly at high conversion rates, the grid-search method achieves slightly higher accuracy (**Figure 4C**). Representative trajectories are shown in (**Figure 4D**). The grid search mapping produces consistently better results than the analytical mapping. Heatmaps spanning a broader range of dominance values, h, are provided in **Supplementary Figure S4**, confirming that these qualitative patterns persist across dominance regimes.

**Figure 4.**
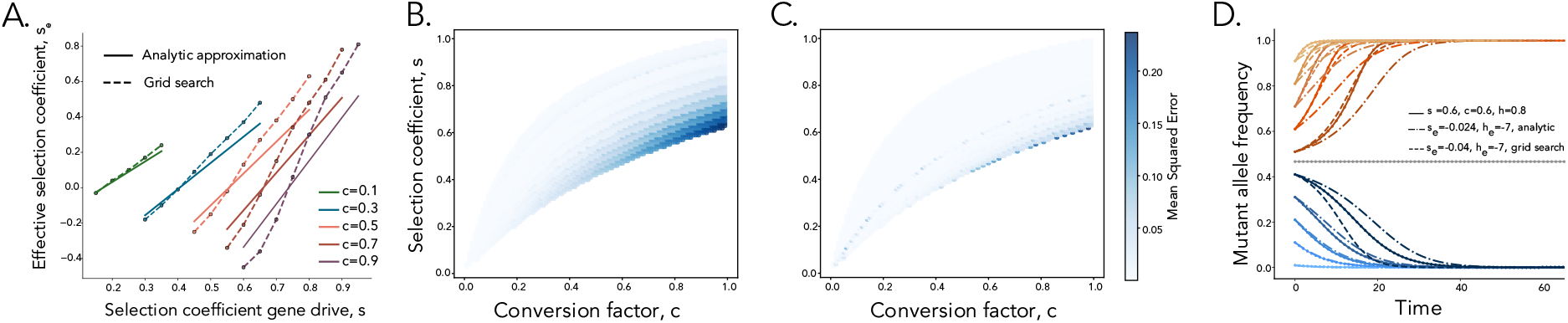
Unstable equilibrium regime: Mapping to a diploid Mendelian model. **A.** Effective diploid selection coefficient (s_e_) as a function of the gene-drive selection coefficient (s) for h = 0.8 and different conversion rates (c). Dash–dot lines denote analytic approximations using eq. (10), and dashed lines show grid-search results. **B, C**. Heatmaps of the mean squared error (MSE) between mapped Mendelian trajectories and the original gene-drive trajectories across (s, c) space, using the analytic mapping (**B**) and the grid-search mapping (**C**). **D**. Representative examples of gene-drive allele frequency trajectories (s = 0.6, c = 0.6, h = 0.8) at different initial allele frequencies. The unstable equilibrium point is q_unstable_ = 0.47. All panels assume dominance h = 0.8.

We also evaluated similar mappings for the stable internal equilibrium and loss regimes (presented in **Supplementary Figure S5**). Overall, the analytic diploid mapping provides a computationally efficient and accurate approximation across all regimes, reliably capturing both equilibrium structure and allele-frequency trajectories. The grid-search mapping yields modest improvements near nonlinear regime boundaries and at extreme parameter values.

### Identifying gene-drive swamping scenarios

We next applied our mapping framework to study *gene swamping*, a containment outcome in which sufficiently high migration can prevent a gene drive from establishing in a connected population, ultimately driving the allele to extinction (21). Our goal was to examine whether gene swamping outcomes can be coherently identified using the mapping and the investigation of the Mendelian model, which would potentially allow expanding the theory of gene swamping in gene drives using results from classic migration-selection theory. To model this process, we used a well-known two-deme migration–selection framework (8, 9, 28, 36). In this formulation, deme 1 represents the *target* habitat where the gene drive is released, and deme 2 represents a connected *non-target* habitat. The drive experiences parameters (s_1_, c_1_, h_1_) in the target deme that place it within the fixation regime, while in the non-target deme it experiences parameters (s_2_, c_2_, h_2_) (**Figure 5A**). The two demes exchange individuals at a symmetric migration rate m. We initialize the drive allele at frequency q_init_ = 0.1 in deme 1, representing a moderate release, and at frequency 0 in deme 2 to reflect its absence in the surrounding population at the time of introduction.

**Figure 5.**
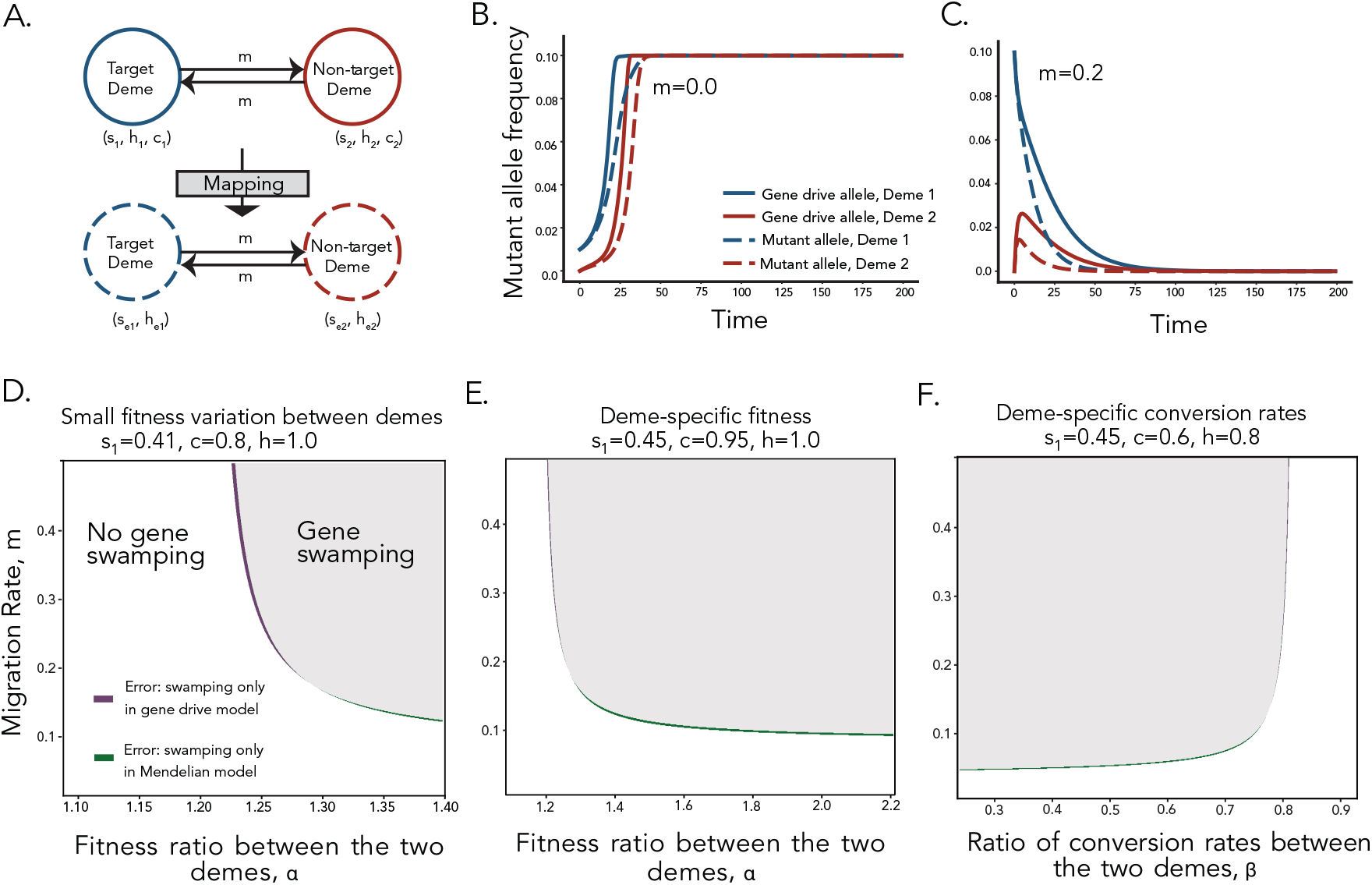
Predicting gene swamping in gene drive deployment. **A.** Mapping in a two-deme model. **B, C**. Representative examples of allele-frequency dynamics in the two demes, with s = 0.45, c = 0.95, and h = 1.0. Solid lines denote gene-drive allele trajectories, and dashed lines denote the corresponding mapped mutant allele trajectories in the Mendelian (non–gene-drive) model. **B**. Gene-drive allele fixes in both demes. Here, migration rate m = 0.05 and *α* = 1.2. **C**. Gene-drive allele is lost in both demes. Here, migration rate m = 0.2 and *α* = 1.4. **D–F**. Gene swamping predictions from the mapped model compared with gene-drive simulations. **D**. Scenario with small differences in fitness costs across environments. Here, s_1_ = 0.41, c = 0.8, and h = 1.0. **E**. Scenario with large, designed differences in fitness costs across environments, Here, s_1_ = 0.45, c = 0.95, and h = 1.0. **F**. Scenario with different conversion mechanisms across environments, Here, s = 0.45, c_1_ = 0.6, and h = 0.8.

To illustrate that gene swamping can arise in this model, we first show that there exist parameter sets in which low and high migration rates lead to drastically different evolutionary outcomes(**Figure 5B, C**). When migration is low (m = 0.05), the gene-drive allele spreads from the target deme into the non-target deme and ultimately fixes in both populations. In contrast, when migration is high (m = 0.2), the continual influx of wild-type individuals from deme 2 suppresses the drive frequency in deme 1, leading to the loss of the drive from both demes. Thus, although the gene-drive parameters in deme 1 would result in fixation in an isolated population, sufficiently strong migration continually pulls the allele frequency toward zero, producing a classic gene-swamping outcome. Notably, the mapped Mendelian model reproduces both the fixation and gene-swamping scenarios almost identically to the full gene drive system, demonstrating that the mapping reliably captures the qualitative dependence of gene-drive dynamics on migration rate (**Figure 5B, C**).

To systematically characterize when gene swamping dynamics can be predicted by our mapping framework, we explore three different scenarios describing how the gene-drive parameters in the non-target deme, (s_2_, c_2_, h_2_), may differ from their values in the target deme. These scenarios capture both naturally occurring environmental differences and engineered containment strategies. In the first scenario, ‘*environmental fitness variation*’, we consider the case where natural environmental variation may alter the fitness cost associated with the gene-drive allele. We model this by allowing the drive to experience a higher fitness cost in the non-target deme, with *α* = s_2_/s_1_ ≈ 1.2 (**Figure 5D**). In the second scenario, ‘*engineered fitness asymmetry*’, we consider the deliberate design of a gene drive that interacts with a particular environmental condition in the target deme (for example, to enable containment of the gene drive) (2, 10, 31, 34). Under this design, the drive may be only mildly deleterious in the target habitat (s_1_ = 0.4) but strongly deleterious elsewhere (s_2_ = 0.9) (**Figure 5E**). In the third scenario, ‘*environment-dependent conversion efficiency*’, we consider the case where environment effects the gene drive conversion efficiency. We model this by assigning high conversion in the target deme (c_1_ = 0.9) and low conversion in the non-target deme (c_2_ = 0.2). The contrast is captured by the parameter *β* = c_2_/c_1_ (**Figure 5F**). Although technically challenging to engineer, such context-dependent conversion mechanisms have been proposed as a pathway to reduce spread into undesired habitats (13, 30).

Across all three scenarios, the mapping to the Mendelian model successfully predicted the qualitative outcomes of the full gene drive simulations, correctly identifying the migration thresholds at which gene swamping occurs. These results highlight the potential of the mapping framework for guiding the design of strategies aimed at preventing spillover or ensuring drive reversal.

Finally, we evaluate a constrained version of the mapping in which the effective dominance coefficient in the non-target deme was forced to equal that of the target deme (h_e,1_ = h_e,2_), with only the effective selection coefficient s_e,2_ allowed to vary. As shown in **Supplementary Figure S6**, this constraint substantially reduced the accuracy of the mapping, emphasizing that both effective parameters (s_e_, h_e_) are essential for faithfully capturing the dynamics of the gene drive.

## Discussion

This work establishes a general framework for translating gene-drive models into equivalent Mendelian models, enabling fast and tractable analysis of their evolutionary outcomes. By recasting biased inheritance in terms of effective Mendelian parameters, such as effective selection and dominance coefficients, we show that engineered gene drive systems can, in many cases, be interpreted and predicted using classical population-genetic theory. This mapping provides conceptual clarity and, importantly, allows researchers to leverage decades of modeling results and analytical tools from Mendelian population genetics to study and predict gene-drive behavior.

We show that, in the fixation regime, a one-parameter haploid mapping provides a surprisingly accurate first-order approximation, capturing the key features of gene-drive spread. Extending the mapping to the diploid case, via effective selection and dominance parameters (s_e_, h_e_), not only improves accuracy across the parameter space but also recovers all four dynamical regimes of the gene-drive model: fixation, loss, stable equilibrium, and unstable threshold equilibrium. This improvement is most pronounced near regime boundaries, where heterozygote effects strongly influence both transient dynamics and long-term outcomes (16, 33).

Across regimes, we apply two complementary mapping strategies: an analytic approximation and a grid-search–based mapping. The analytic formulation offers a closed-form, interpretable, and computationally efficient solution that performs well throughout most of the parameter space. The grid-search mapping, although more computationally demanding, provides higher fidelity in nonlinear boundary regions where small parameter changes produce qualitative shifts in system behavior. This superior performance reflects the capacity of trajectory-based comparisons to accommodate edge effects that challenge local linear approximations. Together, these results demonstrate that gene-drive outcomes can be predicted and interpreted within the framework of standard selection theory.

We demonstrate the power of our approach by applying the analytic diploid mapping to predict gene swamping in a two-deme migration–selection system (18, 21, 28). The mapped Mendelian models accurately reproduced gene drive outcomes across a broad range of migration rates, correctly identifying transitions between fixation and loss of the gene drive allele. Notably, the mapping also captured outcomes in engineered scenarios designed to challenge containment, including environmental variation in fitness costs, deliberate habitat-specific fitness asymmetry, and environment-dependent conversion rates. In all cases, the mapping reproduced the qualitative behavior of full gene drive simulations, underscoring its utility for forecasting confinement, spillover, and system collapse.

While our analysis focuses on deterministic, single-locus dynamics in constant environments, real populations experience demographic stochasticity, fluctuating ecological conditions, resistance evolution, and ecological feedbacks (7, 14, 27). Extending the mapping framework to stochastic models, spatially heterogeneous landscapes, or multilocus gene-drive architectures is therefore an important direction for future work.

Overall, our results demonstrate that key features of gene-drive dynamics can be represented with classical population-genetic models, enabling the use of the extensive theoretical machinery developed for Mendelian systems. By linking non-Mendelian drive systems to standard selection theory, this framework provides both conceptual clarity and a computationally efficient foundation for predicting gene-drive spread and for designing safe, robust, and controllable drive strategies, even under more realistic ecological conditions.

## Methods

### Mapping error calculation

In both the analytical and grid-search-based mappings, we quantify the similarity between the gene drive trajectory q(t), parameterized by the gene drive parameters (s, c, h), and the corresponding q (t), ′computed by the mapped effective parameters (s_e_, h_e_) (or se for haploid Mendelian model) by the mean squared error (MSE) between their allele frequency trajectories:

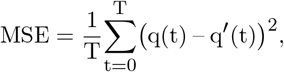

where T denotes the final time step when the gene drive allele frequency reaches a stable equilibrium. This trajectory-level error metric aims to provide a direct comparison of the full temporal dynamics of the two models.

### Grid search

For a gene drive parameter set (s, c, h), the grid search approach finds the best parameter set (s_e_, h_e_) (or s_e_ if we are mapping to the haploid model) by searching through the Mendelian parameter space on a fine grid. We consider s_e_ values in the range [–10, 1] with step size 0.01, and h_e_ values in the range [0, 5] with step size 0.01. In the mapping from gene drive model to diploid Mendelian model, we search both s_e_ and h_e_ for mapping in the fixation regime, but we search only s_e_ for mappings in the unstable or stable regime since h_e_ can be analytically computed from gene drive parameters.

For each candidate Mendelian parameter pair (s_e_, h_e_) (or s_e_), we simulate the deterministic Mendelian model using eq. (4). The Mendelian trajectory is compared with the gene drive trajectory at each discrete time point, and the MSE is computed to quantize the overall difference between allele frequencies across the full trajectory. This ensures that the mapping captures the full transient dynamics of two models. The optimal Mendelian parameters (s_e_, h_e_) are defined as the pair that minimizes the MSE over the grid. Although computationally intensive, this exhaustive grid search provides a robust baseline for mapping gene drive dynamics onto an effective Mendelian framework by explicitly minimizing trajectory-level discrepancies.

## Data availability

Custom scripts were used for the simulations and data analyses. All C++ and Python simulation code is available on Github at https://github.com/MONICAWAN1/Gene_Drive_Mapping. All packages used for analysis and visualization are open-source.

## Acknowledgments

This research was done using resources provided by the Open Science Grid, which is supported by the National Science Foundation award 1148698, and the U.S. Department of Energy’s Office of Science.

## Funding

We gratefully acknowledge support from the National Institutes of Health, National Institute of General Medical Sciences (award no. R35GM147445 to OC), from the National Science Foundation (NSF CAREER Award 2442397 to OC), and from the Israel Science Foundation (ISF award 2049/21 to GG).

## Conflicts of interest

The authors declare no conflicts of interest.

## Notes

### Competing Interest Statement

The authors have declared no competing interest.

