## Supplementary Information for "Mapping Gene Drive Dynamics onto Mendelian Models"

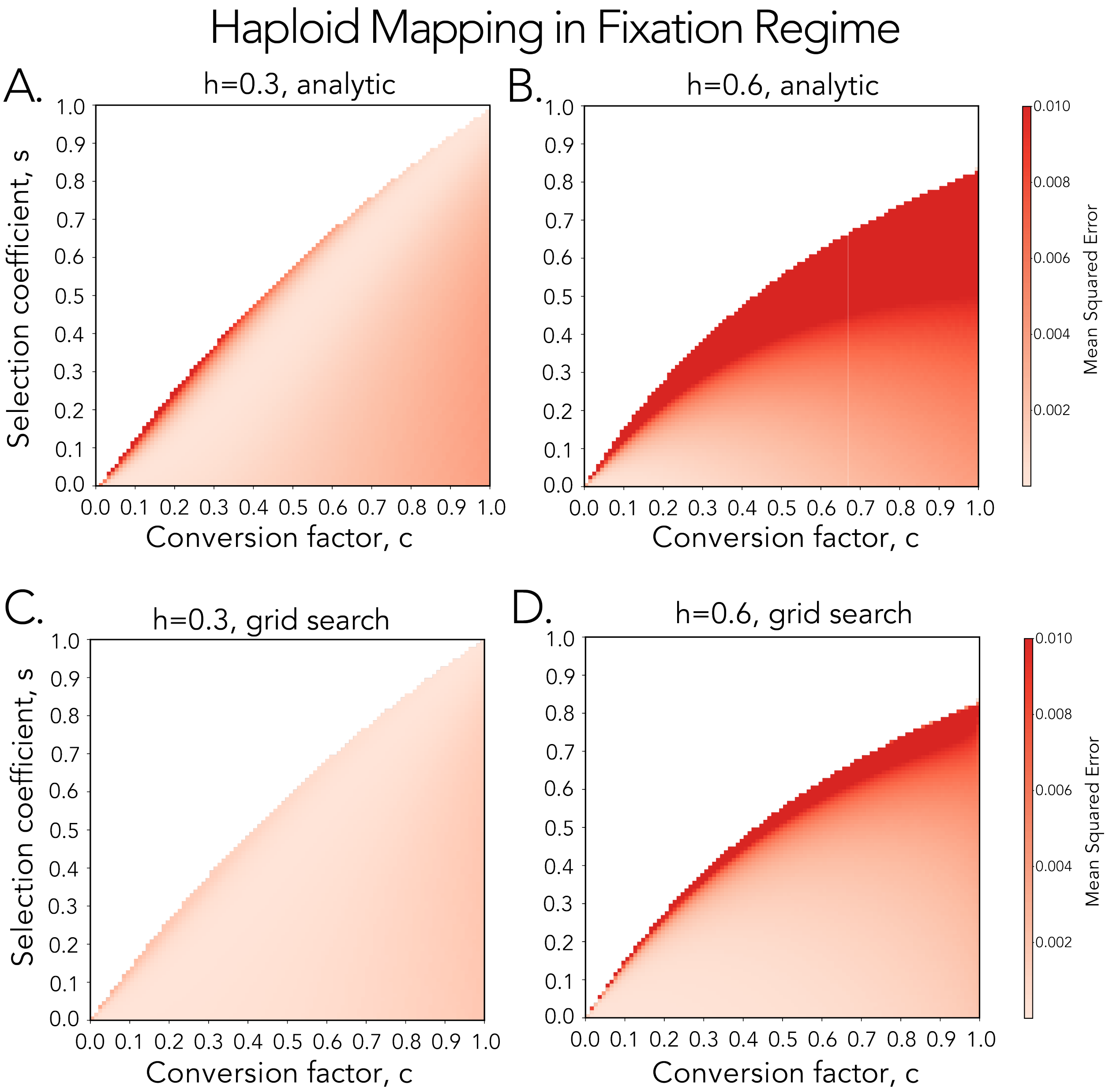


Figure S1: Heatmaps showing the mean squared error (MSE) between the mapped haploid approximations and the original gene-drive trajectories across the $(s, c)$ parameter space, across different values of gene-drive dominance rate, $h$.

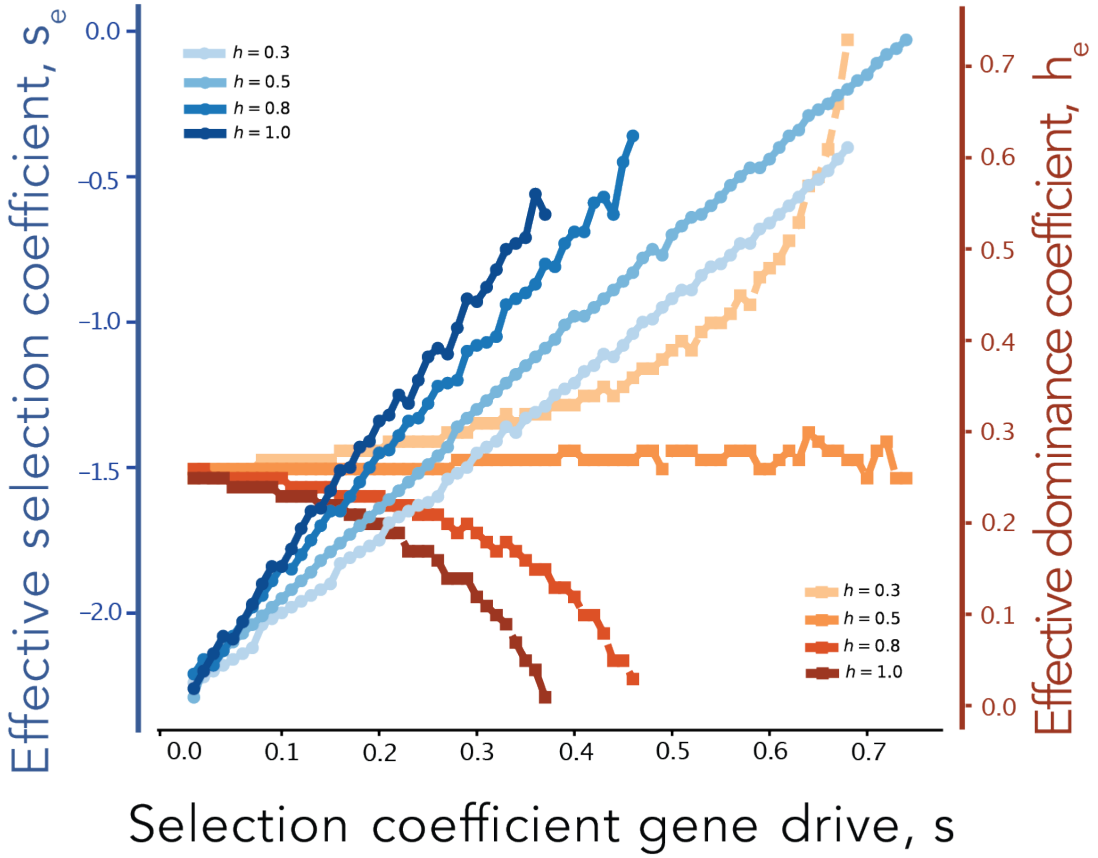


Figure S2: Effective diploid selection coefficients ($s_e$) and effective dominance coefficients ($h_e$) obtained from grid-search mapping, plotted as functions of the gene-drive selection coefficient ($s$) across different values of dominance coefficient ($h$). The conversion rate ($c$) is fixed at 0.6.

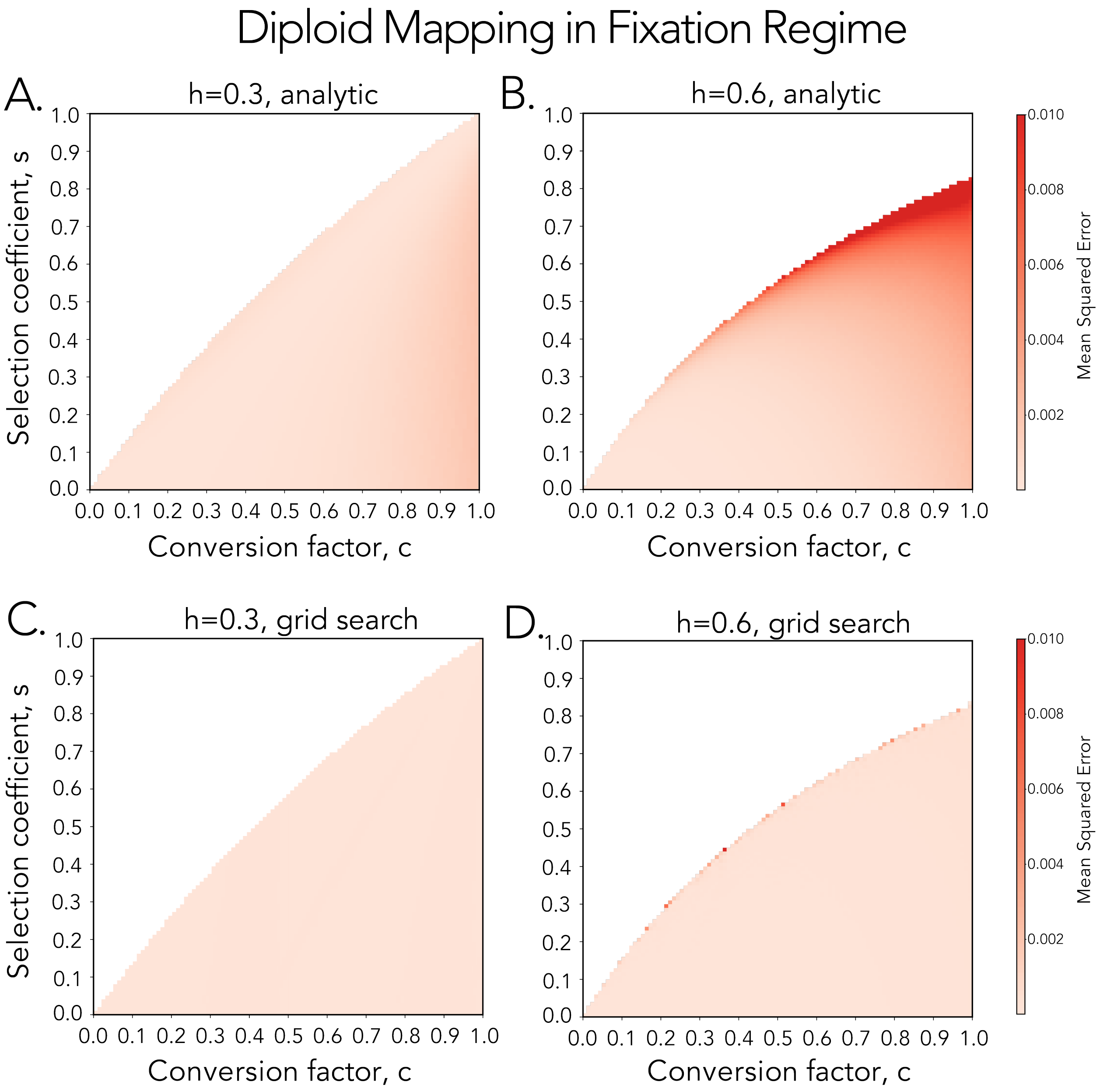


Figure S3: Heatmaps showing the mean squared error (MSE) between the mapped diploid approximations and the original gene-drive trajectories across the $(s, c)$ parameter space observed across different values of gene-drive dominance rate $(h)$. Panels A and C show mapping errors when $h = 0.3$; panels B and D show mapping errors when $h = 0.6$. Panels A and B use the analytic mapping; panels C and D use the grid-search mapping.

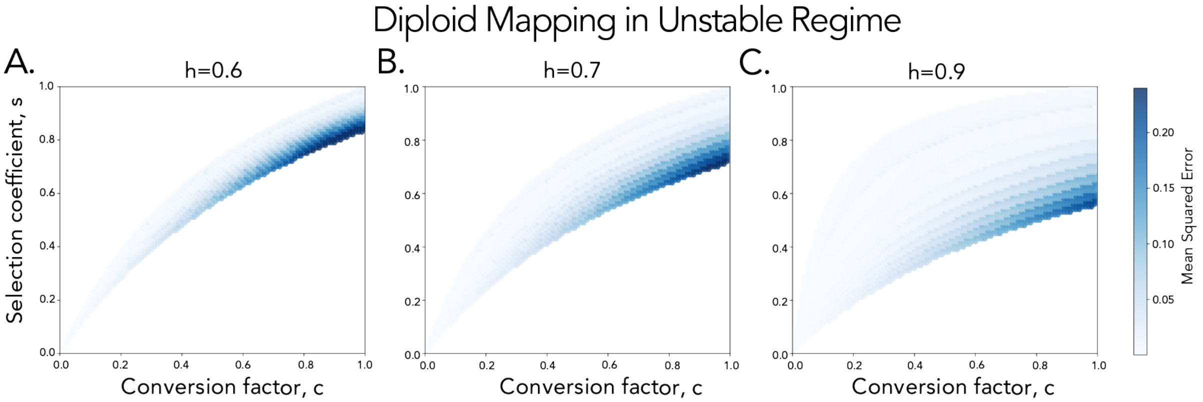


Figure S4: Heatmaps showing the mean squared error (MSE) between the diploid analytic mapping and the original gene-drive trajectories across the $(s, c)$ parameter space in the unstable regime, evaluated across a range of dominance values $h$.

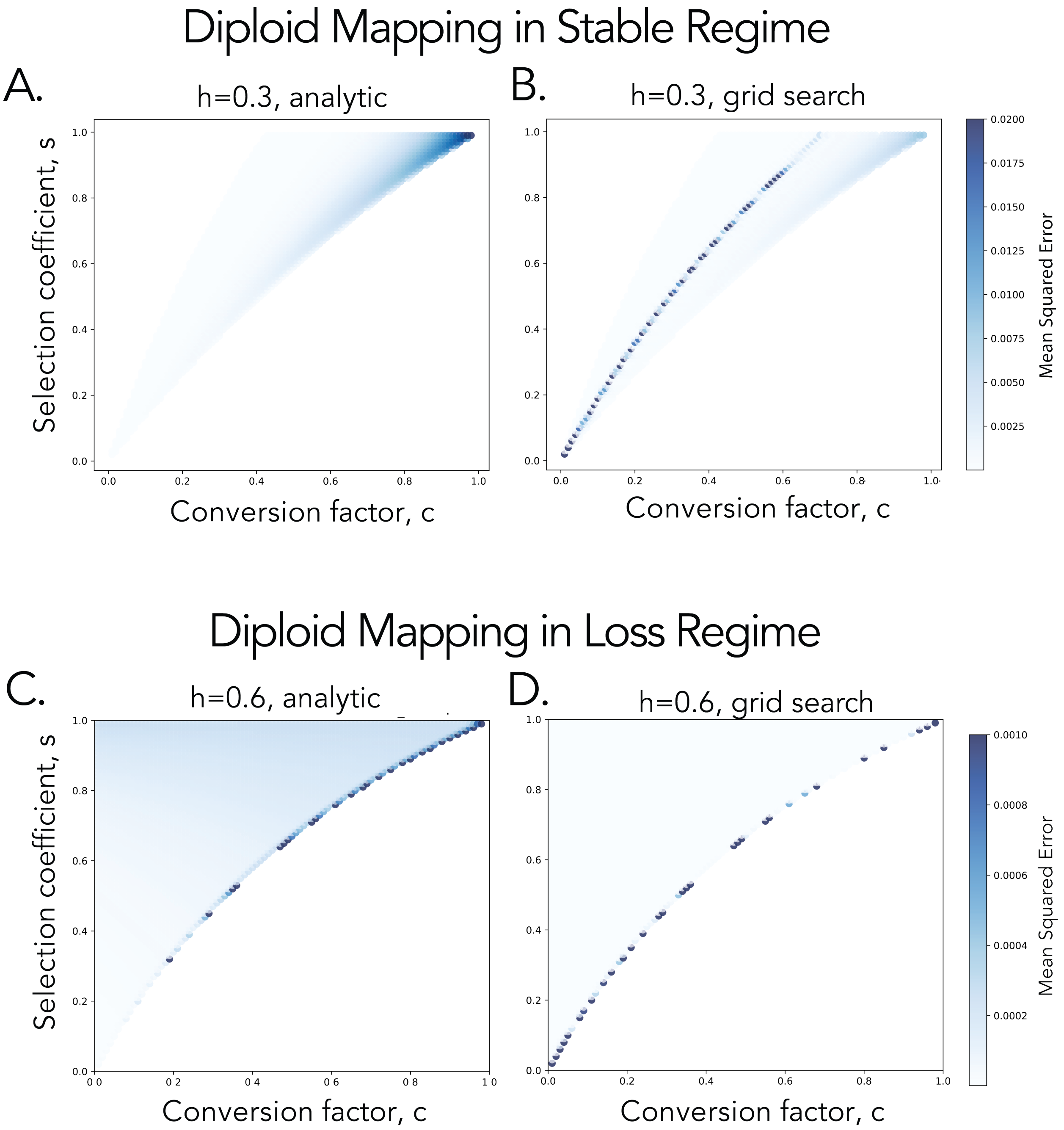


Figure S5: Heatmaps showing the mean squared error (MSE) between the mapped diploid approximations and the original gene-drive trajectories across the $(s, c)$ parameter space. Panels A and B show the stable equilibrium regime. Panels C and D show the loss of gene drive regime.

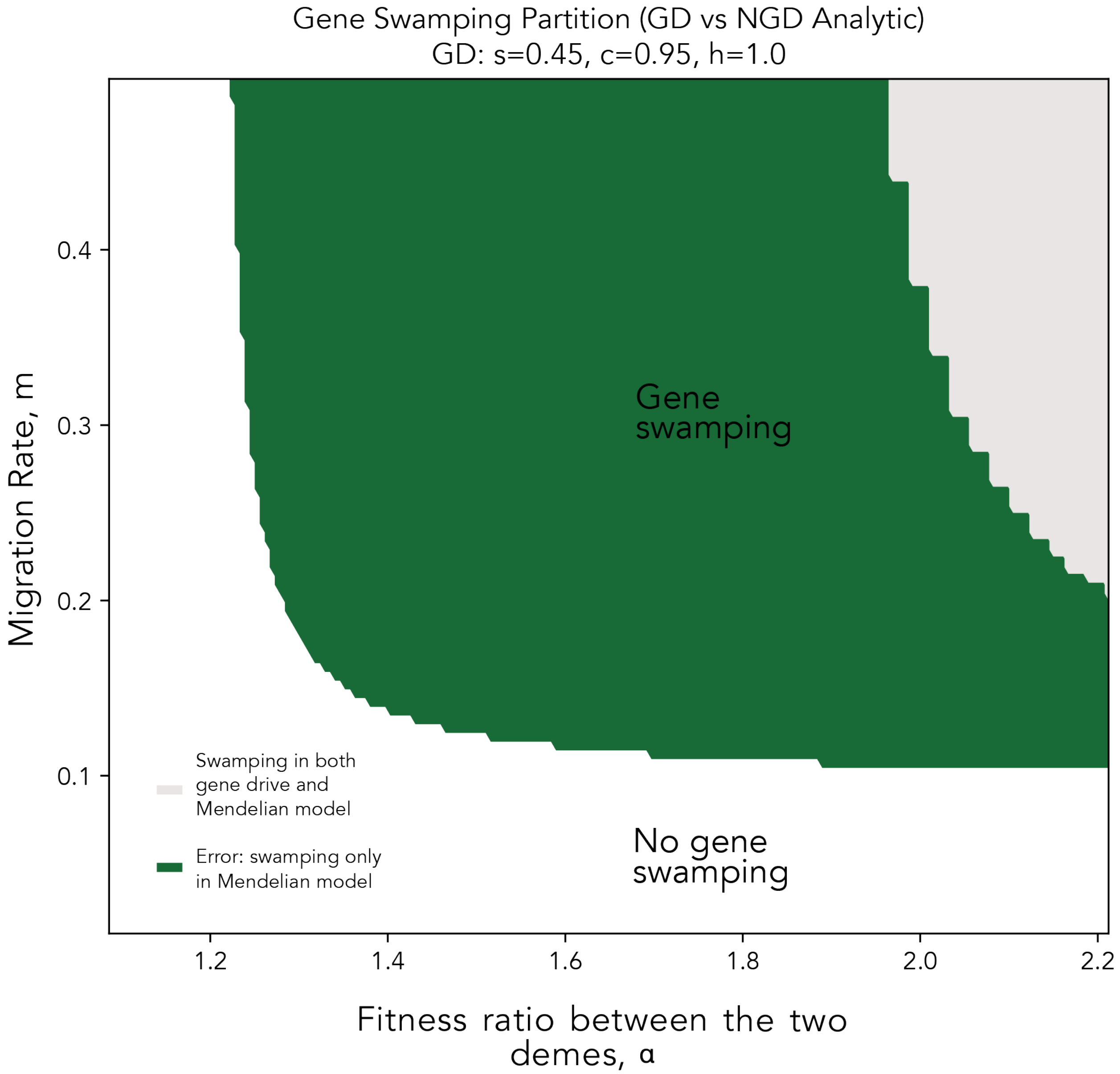


Figure S6: Example scenario in which the mapping is constrained by enforcing $h_{e,2} = h_{e,1}$ and estimating only $s_{e,2}$ in the non-target deme, in order to isolate the effect of differences in $s_e$'s. Under this restriction, the mapped NGD model exhibits a larger deviation from the full gene-drive dynamics.
